# Inference of the Human Polyadenylation Code

**DOI:** 10.1101/130591

**Authors:** Michael K. K. Leung, Andrew Delong, Brendan J. Frey

## Abstract

Processing of transcripts at the 3’-end involves cleavage at a polyadenylation site followed by the addition of a poly(A)-tail. By selecting which polyadenylation site is cleaved, alternative polyadenylation enables genes to produce transcript isoforms with different 3’-ends. To facilitate the identification and treatment of disease-causing mutations that affect polyadenylation and to understand the underlying regulatory processes, a computational model that can accurately predict polyadenylation patterns based on genomic features is desirable. Previous works have focused on identifying candidate polyadenylation sites and classifying sites which may be tissue-specific. What is lacking is a predictive model of the underlying mechanism of site selection, competition, and processing efficiency in a tissue-specific manner. We develop a deep learning model that trains on 3’-end sequencing data and predicts tissue-specific site selection among competing polyadenylation sites in the 3’ untranslated region of the human genome.

Two neural network architectures are evaluated: one built on hand-engineered features, and another that directly learns from the genomic sequence. The hand-engineered features include polyadenylation signals, cis-regulatory elements, n-mer counts, nucleosome occupancy, and RNA-binding protein motifs. The direct-from-sequence model is inferred without prior knowledge on polyadenylation, based on a convolutional neural network trained with genomic sequences surrounding each polyadenylation site as input. Both models are trained using the TensorFlow library.

The proposed polyadenylation code can predict site selection among competing polyadenylation sites in different tissues. Importantly, it does so without relying on evolutionary conservation. The model can distinguish pathogenic from benign variants that appear near annotated polyadenylation sites in ClinVar and inspect the genome to find candidate polyadenylation sites. We also provide an analysis on how different features affect the model’s performance.

## Introduction

Polyadenylation is a pervasive mechanism responsible for regulating mRNA function, stability, localization, and translation efficiency. As much as 70% of human genes are subject to alternative polyadenylation (APA) and wide-spread mechanisms have been found which influence its regulation (1). By selecting which polyadenylation site (PAS) is cleaved, different transcript isoforms that vary either in their coding sequences or in their 3’ untranslated region (3’-UTR) can be produced. Transcripts differentially cleaved can influence how they are regulated. For example, longer variants can harbor additional destabilization elements that alter a transcript’s stability (2), and shortened variants can escape regulation from microRNAs, which have been observed in various cancers (3, 4). Furthermore, APA can be tissue-dependent, so a single gene can generate different transcripts, for instance, based on the tissue in which it is expressed (5). One mechanism of APA regulation occurs at the level of the sequences of the transcript. The presence or absence of certain regulatory elements can influence which PAS is selected. PAS selection is also influenced by its position relative to other sites. A computational model that can accurately predict how polyadenylation is affected by genomic features as well as cellular context is highly desirable to understand this widespread phenomenon. Moreover, several inherited diseases have been linked to errors in 3’-end processing (6). Such model would enable the exploration of the effects of genetic variations on polyadenylation and their implications for disease.

Here, we present the polyadenylation code, a computational model that can predict alternative polyadenylation patterns from transcript sequences. While there have been various previous works in classifying whether a stretch of sequence contains a PAS (7–12), or characterizing whether a PAS is tissue-specific (13, 14), many of them are aimed at improving gene annotations and understanding which features are involved in APA regulation, and does not address the question of predicting how APA sites are variably selected. Here, we tackle this question directly by developing a model that can infer how sites in the same gene are selected for cleavage and polyadenylation. This score, which we refer to as PAS strength, describes the efficiency in which a PAS is recognized by 3’-end processing machinery for cleavage and polyadenylation (15). The ability to predict PAS strength enables this model to generalize to multiple prediction tasks, even though it is not explicitly trained for them. For example, the polyadenylation code can be applied to a gene with multiple PAS to determine the relative transcript isoforms that would be produced, in a tissue-specific manner. The model can predict the consequence of nucleotide substitutions on PAS strength, which can be used to prioritize genetic variants that affects polyadenylation. It can also be scanned across the genome to find potential PAS. We demonstrate examples of these applications in this work.

## Results

### Inferring the Strength of a Polyadenylation Site

The goal of this work is to infer a score that describes the strength of each PAS, or the efficiency in which it is recognized by 3’-end processing machinery. The problem would be straightforward if this target variable is directly measurable. However, current sequencing protocols only provide a measurement of the relative transcript abundance from alternative polyadenylation. Various approaches exist in the literature which attempt to quantify the strength of a PAS. For example, normalized read counts are often used, but quantification can be affected by factors such as sequencing biases, transcript length, and RNA decay (16, 17). Some studies classify PAS strength based on whether a canonical polyadenylation signal or other known sequence elements are present near the PAS (8). We believe a more principled approach to predict a quantitative description of the strength of a PAS is to model it as a hidden variable, and infer it from data. Moreover, the position of a PAS relative to neighboring sites affects its selection. Some biological processes and tissues tend to favor PAS at the distal end, whereas cells under disease states tend to utilize PAS that are more proximal (1). Therefore, the model should account for the distance between neighboring sites during training to appropriately reflect the observed relative abundances. Even though the position of a PAS is modeled during training, a desirable characteristic of the model is that during inference, positional information should be optional. This can be useful in regions of the genome where there are insufficient annotation sources to ascertain the distance to a nearby PAS. This would also enable one to apply this model to any DNA sequence, optionally modify the bases within, and see the predicted effect on polyadenylation regulation at a particular site. Should the user want to see how different PAS influence each other, the model would support that, by applying the model on each PAS separately, and optionally including their position if annotation sources are available, to get a better estimate.

### The Polyadenylation Code

The polyadenylation code is a model that can infer tissue-specific PAS strength scores from sequence, and optionally account for the influence of position if given the context. The code takes as input a sequence of length 200 bases centered on a PAS. We benchmark two variations of the model.

The first model is built on hand-engineered features. The genomic sequence is processed by a feature extraction pipeline, which divides the sequence into 4 regions relative to the PAS (Suppl. S1) (18). Some feature are limited to specific regions, namely the polyadenylation signals in the 5’-5’ and 5’-3’ regions, and hexameric cis-elements defined in (18). Other features are computed in all regions, including counts of RNA-binding protein (RBP) motifs that may be involved in polyadenylation, all possible 1 to 4 n-mers counts, and nucleosome positioning features from (19). The feature vector is mapped to a fully-connected hidden layer of a neural network. We will refer to this model as the Feature-Net.

The second model directly learns from the genomic sequence, using a convolutional neural network (Conv-Net) architecture (20), which can efficiently discover sequence patterns without prior knowledge even when the location of the patterns are unknown. The Conv-Net comprises of tunable motif filters which are free to adapt to the input sequence to optimize the predictive performance of the model. It also contains pooling operations that enables the model to focus on select locations in the input sequence that maximally activate the learned filters.

Both models transform the input sequence into a hidden representation, which is subsequently processed through non-linear activation functions to predict polyadenylation patterns by a fully-connected neural network. The architecture factors predictions into two components: a score that describes the tissue-specific PAS strength, followed by a prediction that reflects the observed relative abundance of transcripts.

To account for positional preference of PAS, the log distance between sites is also an input feature for both models. Given two sites, the proximal (5’) site has a position feature of 0, whereas the distal (3’) site has a position feature that is equal to the logarithm of the distance between the distal and proximal site.

Figure 1 shows a schematic of both models. Parameters of the PAS strength predictor are shared. Separate fully-connected hidden layers are each used to make tissue-specific predictions. The architecture promotes the model to first learn a set of parameters based on the sequence features, via the weights of the connections to a hidden layer in the Feature-Net and filters in the Conv-Net respectively, and then learn a set of tissue-specific parameters that match the observed transcript abundances. These tissue-specific parameters model the cell state, which describes the steady-state environment of the cell, such as the protein concentrations in the cytosol, that can affect transcriptional modifications. We do not explicitly define what exactly these cell state parameters consist of or how they factor in the predictions, but rather simply model them as hidden variables and learn them from data. A similar approach has been described in the splicing regulatory code by Xiong et al. (21).

Seven distinct tissue are available in the dataset used to train the model, although there are two different sets of sequencing reads for the naïve B-cells obtained from two different donors (22). We modeled the two naïve B-cell datasets as distinct tissues, and so the model has eight polyadenylation strength prediction outputs, one each tissue. For both models, we choose not to rely on evolutionary conservation to force the model to learn patterns from the genome itself (23), and to decouple the dependence on external conservation tracks, which can limit the variety of the inputs that the model can process.

The polyadenylation code is trained by modeling the relative strength between pairs of competing sites. Each training example consists of two PAS from the same gene, and requires the model to predict the probability that each site would be selected for cleavage and polyadenylation. A softmax function is used to squash the real-value strength predictions into a normalized score that can be interpreted as the probability that one PAS is chosen over another. This score is penalized against training targets of the relative abundances of transcripts for these PAS, which is observed from the sequencing experiment. Most of the results presented in this work are based on the predictions from the PAS strength predictor (i.e. the logits) instead of the relative strength predictions that follows the softmax. Note that the polyadenylation code is trained only to the task of modeling competing site selection. All the predictions in this work are evaluated without any additional task-specific training whatsoever to demonstrate the general applicability of this model.

**Fig. 1:**
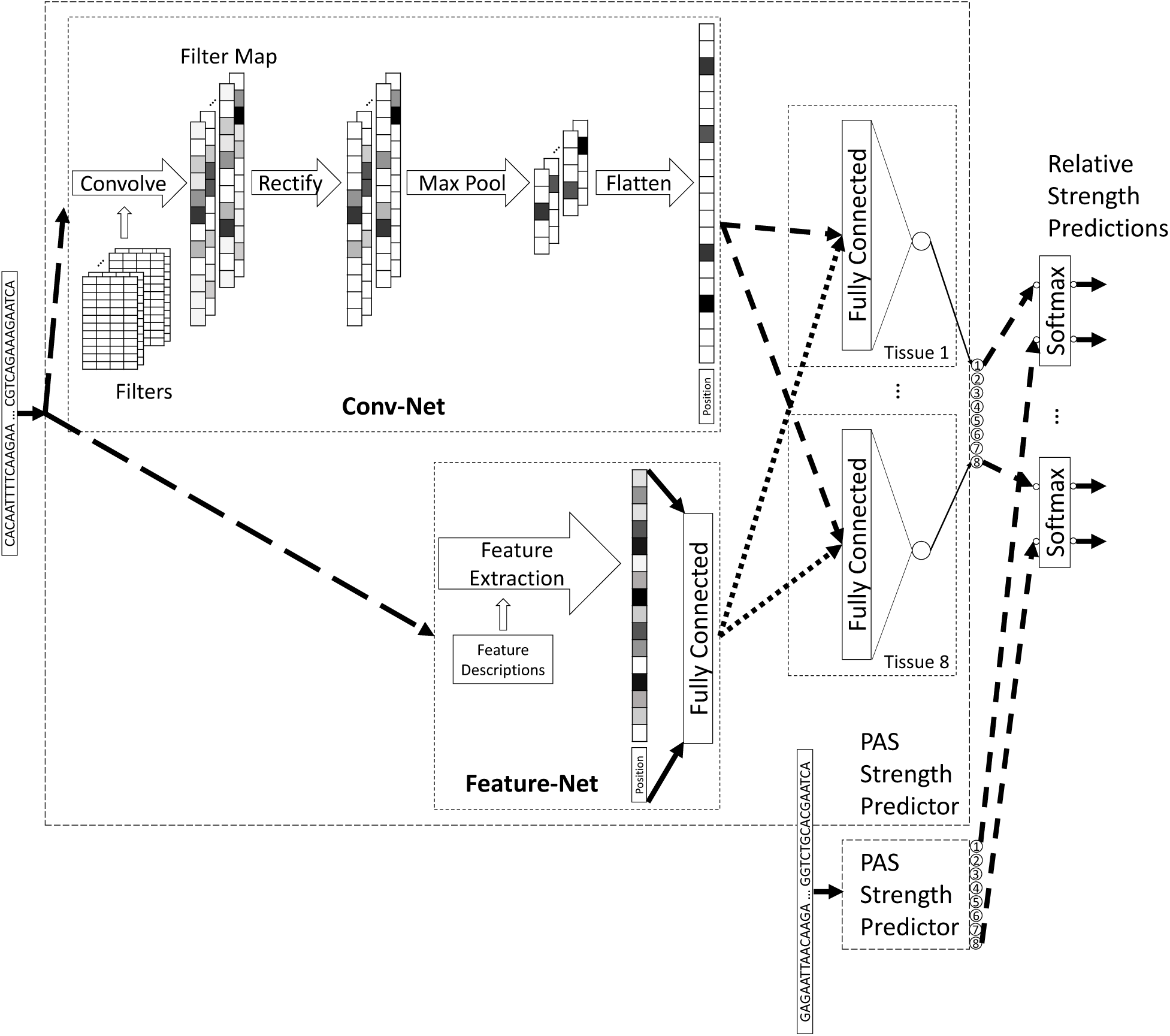
A schematic of the components of the neural network that forms the polyadenylation code. The Conv-Net and the Feature-Net are trained independently. The genomic sequence surrounding each polyadenylation site serves as input to the Strength Predictor. Detailed description on the operations of the Conv-Net can be found in (24).

### Polyadenylation Site Selection

The performance of the polyadenylation code to predict the likelihood that a PAS is selected for cleavage and polyadenylation against a competing site in the same gene is shown in Table 1. These are the tissue-specific relative strength predictions from the model for pairs of PAS, as illustrated in Figure 1. Performance is assessed using the area under the receiver-operator characteristic (ROC) curve (AUC) metric on held-out test data. To compare the code’s performance against a baseline, we also trained a logistic regression (LR) classifier, which is essentially the Feature-Net with hidden layers removed. Predictions from the model based on the convolutional neural network architecture is consistently the best performer. There is sizable performance gain from using the neural network models compared to a logistic regression classifier.

For the more general task of predicting which PAS would be selected in a gene with multiple sites, the polyadenylation code is applied to each PAS in the 3’-UTR region of each gene. The score computed from each site of a gene is used in a classification task to select which PAS is most likely to be selected, the target of which is determined by the site which has the most observed reads in the sequencing data. The metric we report here is the prediction accuracy, or the percentage of genes in which the model has correctly predicted the PAS that has the most observed reads. This is shown in Table 2 for genes with two to six sites, averaged across all tissues.

**Table 1:** PAS Selection Performance Between Competing Sites

| Tissue Type | AUC |  |  |
| --- | --- | --- | --- |
|  | LR | Feature-Net | Conv-Net |
| Brain | 0.826 ± 0.010 | 0.869 ± 0.007 | 0.895 ± 0.005 |
| Breast | 0.825 ± 0.006 | 0.862 ± 0.003 | 0.886 ± 0.004 |
| ES Cells | 0.849 ± 0.006 | 0.898 ± 0.002 | 0.911 ± 0.006 |
| Ovary | 0.830 ± 0.009 | 0.873 ± 0.006 | 0.895 ± 0.003 |
| Skeletal Muscle | 0.828 ± 0.006 | 0.872 ± 0.005 | 0.893 ± 0.004 |
| Testis | 0.787 ± 0.007 | 0.828 ± 0.005 | 0.856 ± 0.007 |
| B Cells 1 | 0.838 ± 0.005 | 0.880 ± 0.005 | 0.896 ± 0.004 |
| B Cells 2 | 0.832 ± 0.004 | 0.880 ± 0.008 | 0.893 ± 0.007 |
| All | 0.824 ± 0.005 | 0.866 ± 0.004 | 0.889 ± 0.003 |

**Table 2:** PAS Selection Performance in Genes with 2 to 6 Sites

| Number of Sites | Accuracy (%) |  |  |
| --- | --- | --- | --- |
|  | LR | Feature-Net | Conv-Net |
| 2 | 79.6 | 82.5 | 83.5 |
| 3 | 68.3 | 73.0 | 75.5 |
| 4 | 58.9 | 64.4 | 69.8 |
| 5 | 55.6 | 62.8 | 64.0 |
| 6 | 48.5 | 56.4 | 59.7 |

### Pathogenicity Prediction

The advantage of the polyadenylation code is that the inferred PAS strength model can provide a characterization of individual sites based on sequence context. We evaluate whether this model can be used for pathogenicity predictions. The basic approach involves applying the polyadenylation code to the 200 nucleotides sequence associated with a PAS from the reference genome to get a prediction, and then performing another prediction when one or more nucleotides in the sequence is altered. A difference is then computed between the reference and variant prediction. Since there are eight predictions, one for each tissue, we take the largest difference as the score to evaluate pathogenicity. A similar approach has been applied to splicing variants (21). The postulate is that if a variant causes a large change to the strength of a PAS, this can change the relative abundance of differentially 3’-UTR terminated transcripts that deviates from normal, potentially indicating disease associations.

To evaluate the efficacy of this approach, we extracted variants that overlap with our PAS atlas (within 100 bases on either side of the annotated PAS) from the ClinVar database (25). Some of these variants overlap with the terminal exon (e.g. missense mutations) and are manually removed. There are 12 variants that are labeled as pathogenic (CLNSIG=5) and 48 that are labeled as benign (CLNSIG=2) (Suppl. S2). Figure 2 shows the ROC curve for this classification task.

The polyadenylation code can predict pathogenic variants from benign ones with an AUC of 0.98 ± 0.02 and 0.97 ± 0.02, for the Conv-Net and Feature-Net respectively, both with a p-value of < 1 × 10-8. Even though the AUC’s are essentially identical for both models, there is clear advantage in the performance characteristic of the Conv-Net: it outperforms in the low false positive rate region where variant classification matters. For these predictions, we used an input of zero for the positional feature of the strength model, since each variant is not analyzed with respect to neighboring sites. However, in general, it may be advantageous to incorporate this information. For example, a variant may cause a large change in a nearby PAS, but if there is a much stronger neighboring PAS in the same gene, the effects of the variant may be dwarfed by this neighbor, and therefore not have any significant mechanistic effects.

We further evaluated how the code compares with four phylogenetic conservation scoring methods: Genomic Evolutionary Rate Profiling (GERP) (26), phastCons (27), phyloP (28), and the 46 species multiple alignment track from the UCSC browser (29). We also compare the scores with Combined Annotation-Dependent Depletion (CADD), a tool which scores the deleteriousness of variants (30). Overall, as shown in Figure 2, the pathogenicity score from the polyadenylation code compares favorably, even though it has not been explicitly trained for this task. It is worth noting that although the polyadenylation code performed well for this ClinVar dataset, in general, a large difference in PAS strength does not necessarily imply pathogenicity, which is a phenotype that can be many steps downstream of 3’ -end processing (31).

The model can also be used to search for potential variants that would affect the regulation of polyadenylation. To visualize this approach, we applied the model and generated a mutation map (24) to a 100 nucleotide sequence in the human genome, where ClinVar mutations associated with β-thalassemia which affect the polyadenylation signal are present (32). As shown in Figure 3, the polyadenylation signal is identified as an important region relative to other bases in the sequence.

**Fig. 2:**
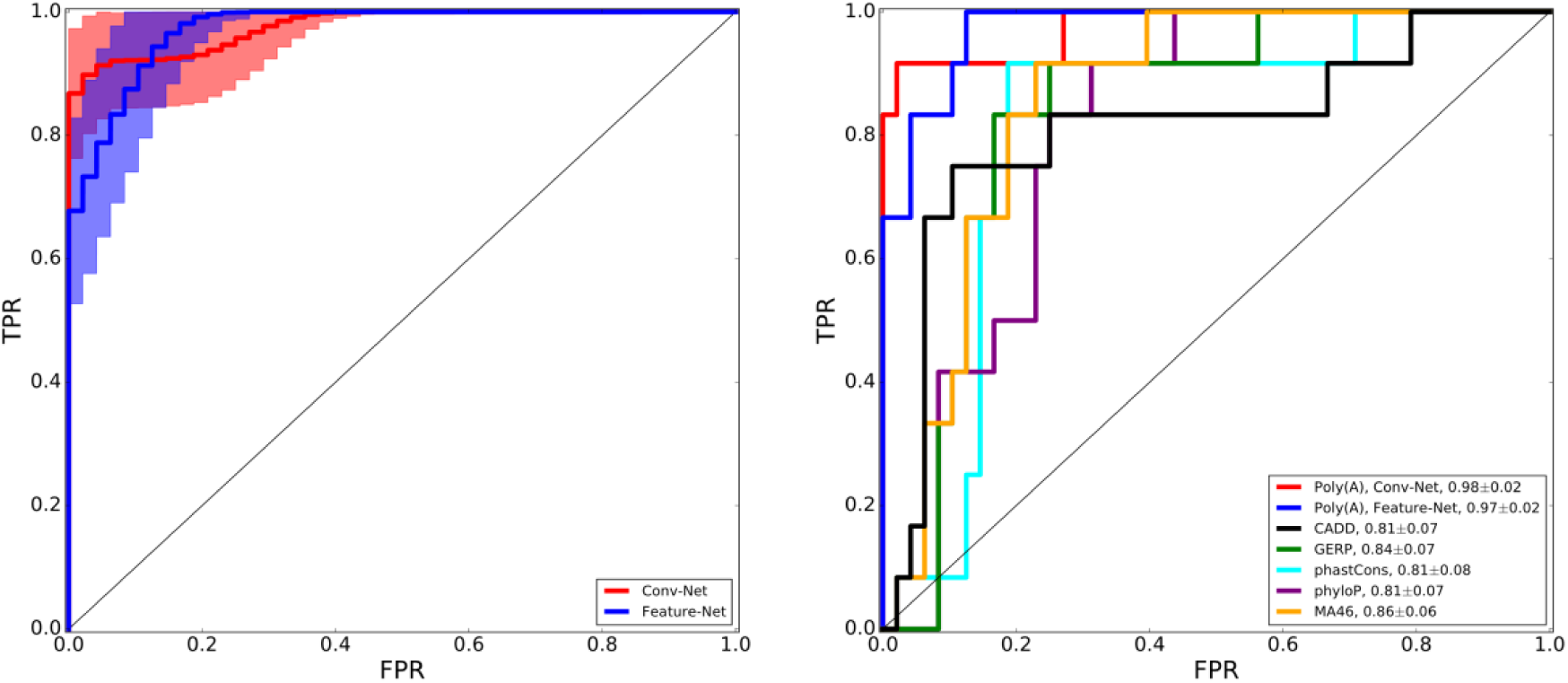
Classification performance on ClinVar variants near polyadenylation sites. (left) ROC curves of variant classification using the polyadenylation code. The shaded region shows the one standard deviation zone computed by bootstrapping. (right) ROC curves comparing the polyadenylation code’s performance against other predictors. AUC values are shown in the figure legend.

**Fig. 3:**
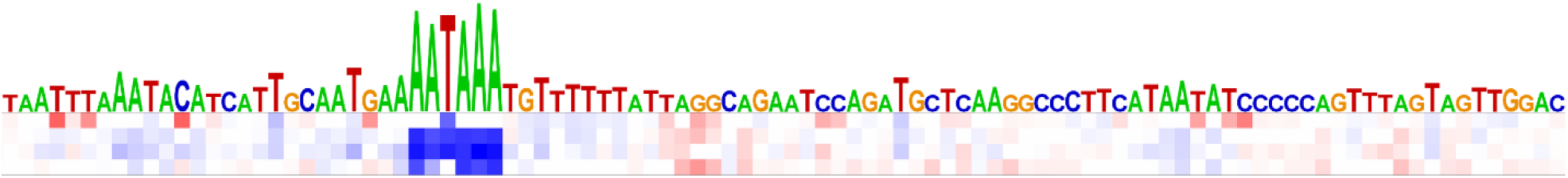
A mutation map of the genomic region chr11: 5,246,678-5,246,777. Each square represents a change in the score if the original base is substituted. The substituted base is represented in each row in the order ‘ACGT’. Red/blue denote a mutation that would increase/decrease the likelihood of the PAS for cleavage and polyadenylation.

### Polyadenylation Site Discovery

The polyadenylation code is trained by centering the input sequence around a PAS. As the PAS is translated away from the center of the 200 nucleotides input sequence, or when a PAS is absent, it stands to reason that the predicted strength of the sequence would be reduced, due to the lack of sequence elements necessary for cleavage and polyadenylation. Naturally, we asked whether the PAS strength predictor can be translated across the genome to find potential PAS. While there have been previous works on this task (7–9), our model is not explicitly trained for this.

Suppl. S3 shows an example of a predicted PAS track across a section of the human genome by applying the Conv-Net strength model in a base-by-base manner. The average strength prediction from all eight tissue models, without application of any filtering or thresholding, is shown. For this example, we chose a region of the genome with multiple PAS, and where there are differences between annotation sources.

The set of predicted peaks labeled region A are present in all annotation sources. It is not a single sharp peak, indicating that various PAS are possible in that region. This agrees with the GENCODE Poly(A) track, which indicates that there are two peaks in this region, as well as 3’-Seq, which shows that there are RNA-Seq reads that map across a broad region for various tissues. As mentioned earlier, the precise location for cleavage and polyadenylation is not exact. Region B is less well-defined, is weaker, and approximately aligns with the predicted positions from another PAS predictor (7), as well as the muscle track from PolyA-Seq (in light gray). Finally, a small peak is observed in Region C, predicted to be a very weak PAS, which is present in PolyA-Seq. Note that the polyadenylation code is trained only from 3’-Seq reads and has no knowledge of RNA-Seq reads from other datasets or other annotation sources.

To assess the polyadenylation code’s ability in discovering polyadenylation site, we created a dataset with positive and negative examples to assess its classification performance. There are no general consensus from previous work on what constitutes a proper criteria on negative sequences or a standardized dataset for this task (33). We therefore defined the evaluation dataset based on our annotations and evidence from 3’-Seq. Positive targets consist of annotated PAS in the 3’-UTR that has 10 or more reads. Since it is generally not appropriate to simply use random genomic sequences or locations, for the negative set, we extracted the two immediately adjacent genomic regions near a PAS to ensure the both the negative and positive sequences have similar compositions (Suppl. S4). The sequences are fed as input into the strength predictor, and the output scores are used for classification. Positional information of the sequences is not used (i.e. all model inputs have a positional feature of zero). Table 3 shows the code’s performance for this task.

**Table 3:**
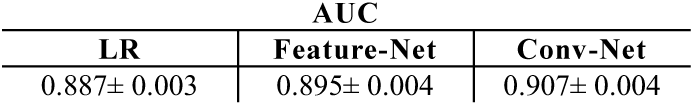
Performance Distinguishing PAS from non-PAS Regions in the 3’-UTR AUC

It is interesting to observe that there is a relatively smaller difference in AUC’s for all models, especially between the Conv-Net and the logistic regression model, compared to previous tasks, which differed more drastically in performance. This is likely because identification of PAS from the genome is a comparatively simple task compared to quantifying its strength, which requires more elaborate integration of genomic context information.

## Discussion

Regulation of polyadenylation is a crucial step in gene expression, and mutations in DNA elements that control polyadenylation can lead to diseases. Accurate, predictive models of polyadenylation will enable a deeper understanding of gene regulation and provide an important new approach to detecting and treating damaging genetic variations. We have presented here the polyadenylation code, a versatile model that can predict alternative polyadenylation patterns from transcript sequences and generalizes to multiple tasks that it was not trained on. Beyond its original trained usage to predict PAS selection from competing sites, it exceeds at classifying variants near PAS and can be used for PAS discovery. In the following sections, we explore properties of both the Feature-Net and Conv-Net models.

### Hand-Engineered Features’ Effect on Model Performance

To understand how different features contribute to the performance of the code, we train models using only individual feature groups. Table 4 shows their classification performance. Even though the polyadenylation signals are generally considered to be a main signature of PAS, they only account for a fraction of the predictive performance for PAS selection compared to the full feature set. Overall, n-mers features are major contributors to the Feature-Net’s performance, which is sufficiently rich to capture many motif patterns. It should be noted that each feature group has a different number of features (see Suppl. S1), and therefore individual features in the larger feature groups may contribute only weakly, but as a whole affect predictions considerably. Position alone have very poor predictive capability, even though it was suggested as being a key feature in determining whether a PAS is used for tissue-specific regulation (14).

To see the contributions of individual features, we computed the gradient of the output with respect to the input feature vector of the neural network. This is referred to as the feature saliency of a prediction, and the gradients of features with large magnitudes can be interpreted as those that need to change the least to affect the prediction the most (34). For this, we computed the feature saliency of all sites in our test set, and selected the features that on average has the largest magnitude. Table 5 shows the top 15 features computed using this method.

**Table 4:** Feature-Net PAS Selection Performance Between Competing Sites w ith Feature Subset

| Feature Group | AUC |
| --- | --- |
| All | 0.866 ± 0.004 |
| Poly(A) Signal | 0.728 ± 0.004 |
| Position | 0.553 ± 0.004 |
| Cis-Elements | 0.608 ± 0.009 |
| RBP Motifs | 0.676 ± 0.009 |
| Nucleosome Occupancy | 0.656 ± 0.006 |
| 1-Mers | 0.762 ± 0.004 |
| 2-Mers | 0.794 ± 0.002 |
| 3-Mers | 0.817 ± 0.004 |
| 4-Mers | 0.833 ± 0.005 |

**Table 5:** Top 15 Features of the Feature-Net

| Rank | Region | Feature Name |
| --- | --- | --- |
| 1 | 5'-3' | Polyadenylation Signal, AAUAAA |
| 2 | --- | Log distance between PAS |
| 3 | 5'-3' | Polyadenylation Signal, AUUAAA |
| 4<br>to<br>15 | 5'-3' | 1-mer, C |
|  | 5'-3' | 1-mer, U |
|  | 5'-3' | 2-mer, AG |
|  | 3'-5' | 2-mer, CA |
|  | 3'-5' | 3-mer, AAA |
|  | 5'-3' | 3-mer, UGU |
|  | 5'-5' | 3-mer, UGU |
|  | 3'-5' | 4-mer, AAAA |
|  | 5'-5' | Cleavage Factor Im, UGUA |
|  | 5'-3' | Polyadenylation Signal, CAAUAA |
|  | 5'-3' | Polyadenylation Signal, AUAAAG |
|  | 5'-5' | Polyadenylation Signal, AGUAAA |

The top three features are consistent for all tissue types. Other features vary slightly between tissues and are grouped together unordered. As expected, the two most common canonical polyadenylation signals are the top features. The log distance between PAS is also deemed to be important. Other features in this list are consistent with mechanisms of core elements known to be involved in cleavage and polyadenylation, including the upstream UGUA motif which the cleavage factor Im complex binds to, and a GU-rich downstream sequence near the polyadenylation site (35). Interestingly, the genomic context upstream of the PAS appears to be more important, as most of the top features are in either the 5’-5’ and 5’-3’ region.

### Convolution Neural Network to Predict the Effect of Genomic Variations

The convolutional neural network doesn’t offer the ability to explore how individual features contribute to the model’s predictions. However, its performance in the tasks that have been evaluated in this work is consistently better than the model with hand-engineered features, and it does so without prior knowledge on polyadenylation. This is surprising at first, but perhaps not so if viewed in the context of other applications of machine learning like computer vision, where hand-engineered features have been largely superseded by models which learn directly from image pixels (36).

On top of this, the Conv-Net has additional advantages that are not available in the Feature-Net. For instance, it is completely free to discover novel sequence elements that may be relevant for polyadenylation regulation from data. An example set of filters from the Conv-Net model is shown in Suppl. S5. It also has the potential to be more computationally efficient. Feature extraction from sequences can be the most computational intensive aspect of a model during inference. This is not required for models that directly operate on sequences. There are additional operations that are required in the Conv-Net, but these computations can be significantly sped up by graphics processing units, which can be important for application of the model to entire genomes.

Since the Conv-Net operates directly on the raw genomic sequence, it also enables one to perform analysis at the single-base resolution more naturally. In the Feature-Net, many features are derived in discrete sections of the genome (four in this case) to reduce the dimensionality of the input. The Conv-Net on the other hand, is more efficient at sharing model parameters, enabling the motif filters to be applied at much finer spatial steps across a genomic sequence (a stride of 1 is used, see S6), while still make overfitting manageable during training.

## Materials and Methods

### Assembling the Polyadenylation Site Atlas

Analysis of human polyadenylation events is confined to the 3’-UTR, where PAS are most frequently located. First, 3’-UTR annotations from UCSC, GENCODE, RefSeq, and Ensembl are combined, where overlapping regions are merged, and each 3’-UTR segment is further extended by 500 bases to capture potential uncharacterized regions. Then, to generate a comprehensive atlas of PAS, polyadenylation annotations and mapped reads from sequencing experiments are inspected to generate an atlas of human PAS in the 3’-UTR. The polyadenylation annotations used include PolyA_DB 2 (37), GENCODE (38), and APADB (39). Mapped reads from PolyA-Seq (40) and 3’-Seq (22) are also analyzed to expand the repertoire of PAS. PAS from different sources overlap, but some sites can be unique to one study due to the differences in cell lines or tissue types as well as sequencing protocol. Due to the inexact nature of 3’-end processing (41), PAS that are within 50 bases of each other are clustered, and the resulting peak marked as the location of the PAS. The final PAS atlas contains 19,316 3’-UTR regions with two or more PAS from genes in the hg19 assembly.

### Quantifying Relative Polyadenylation Site Usage

The polyadenylation code is trained from the relative abundance of transcripts from a 3’-end sequencing experiment of seven distinct human tissues, including the brain, breast, embryonic stem (ES) cells, ovary, skeletal muscle, testis, and naïve B cells from (22). The dataset also contains cell lines, but they are not used. The version of aligned reads which have been processed through the studies’ computational pipeline is used, which include removal of internally primed and antisense reads, as well as application of minimum expression requirements to reduce sequencing noise.

To quantify the relative PAS usage for each gene as the target to train the polyadenylation code, we adopted the Beta model derived from Bayesian inference described in (42), which treats the percent read counts of one site relative to another site as the parameter of a Bernoulli distribution. With this model, the relative PAS usage of one site relative to another, referred to as Φ, is:

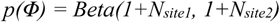

where *N*_*site1*_ and *N*_*site2*_ are the number of reads from two different sites. We use the mean of this distribution as the target to train the polyadenylation code, that is, the PAS usage of site 1 relative to site 2 is (1 + N_site1_) / (2 + N_site1_ + N_site2_). For 3’-UTR regions with more than 2 PAS, we generate pairs of training targets and quantify the reads as above. The assumption is that the relative strength of neighboring PAS can be described by the relative read counts at those sites, even if there are other sites present in the same gene. This assumption simplifies the architecture of the computational model and quantification of relative strength between sites.

### Training the Neural Networks

The polyadenylation code is constructed and trained in Python using the TensorFlow library (43, 44). All hidden units of the neural network consists rectified linear activation units (45). For the Feature-Net, the feature vectors are normalized with mean zero and standard deviation of one. For the Conv-Net, the input uses a one-hot encoding representation for each of the 4 nucleotides. For a sequence of length *n*, the dimension of the input would be 4 × *n*. Padding is inserted at both ends of the sequence so that the motif filters can appropriately scan along the whole length of the sequence. For a motif filter of length *m*, the additional padding on each side of the sequence would be 4 × (*m* - 1), where these additional padding would be filled with the value 0.25, equivalent to an N nucleotide in IUPAC notation. This is similar to what is done in (24).

The parameters of the neural network are initialized according to (46), and trained with stochastic gradient descent with momentum and dropout (47). Predictions from each softmax output are penalized by the cross-entropy function, and its sum across all tissue types are backpropagated to update the parameters of the neural network. Training and testing of the model is performed in a similar fashion as described in (48). Briefly, data is split into approximately five equal folds at random for cross validation. Each fold contains a unique set of genes that are not found in any of the other folds. Three of the folds are used for training, one is used for validation, and one is held out for testing. By selecting which fold is held out for testing, five models are trained. The prediction of these five models on their corresponding test set are used for performance assessment, as well as to estimate variances, for all the tasks analyzed in this work.

The validation set is used for hyperparameters selection. The hyperparameters can be found in Suppl. S6. A graphics processing unit is used to accelerate training and hyperparameter selection by randomly sampling the hyperparameter space. The number of epoch is fixed to 50. Only polyadenylation events with greater than 10 reads are used to train and test the model.

## Supporting Information

**S1.**
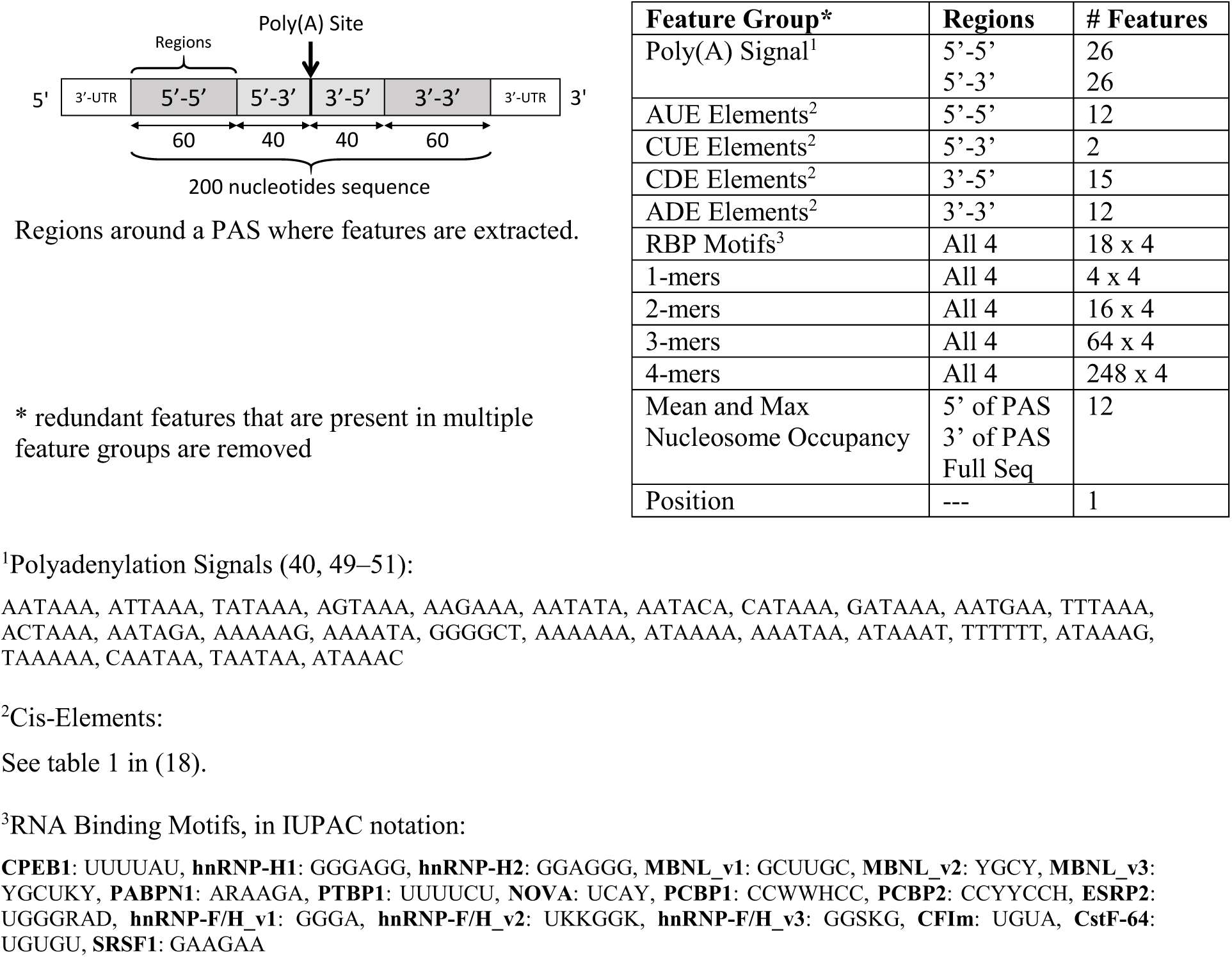
Feature Description of the Feature-Net

**S2.**
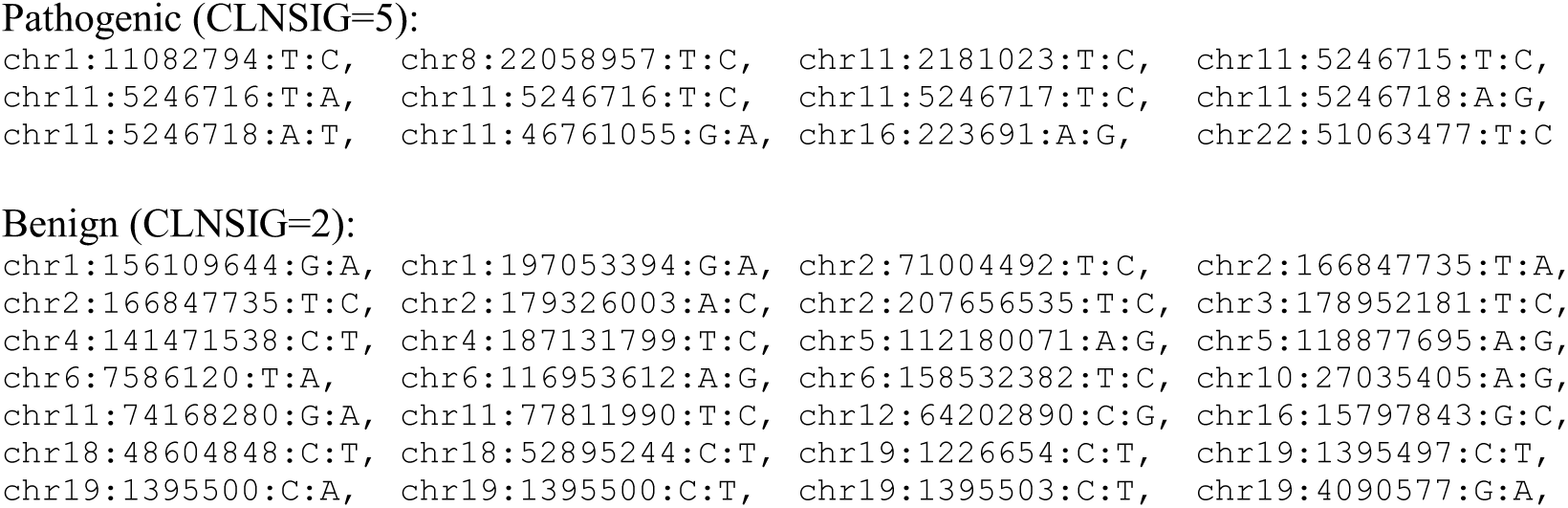
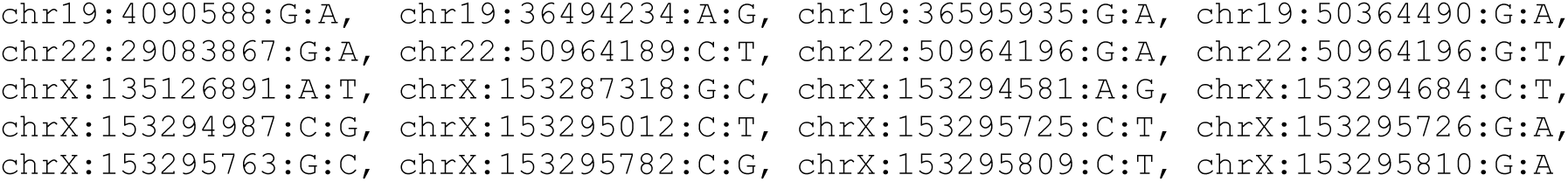
Variants Near Polyadenylation Sites Extracted from ClinVar. Variants are given in notation *chromosome*:*position*:*reference*:*variant*, based on the hg19 assembly.

**S3.**
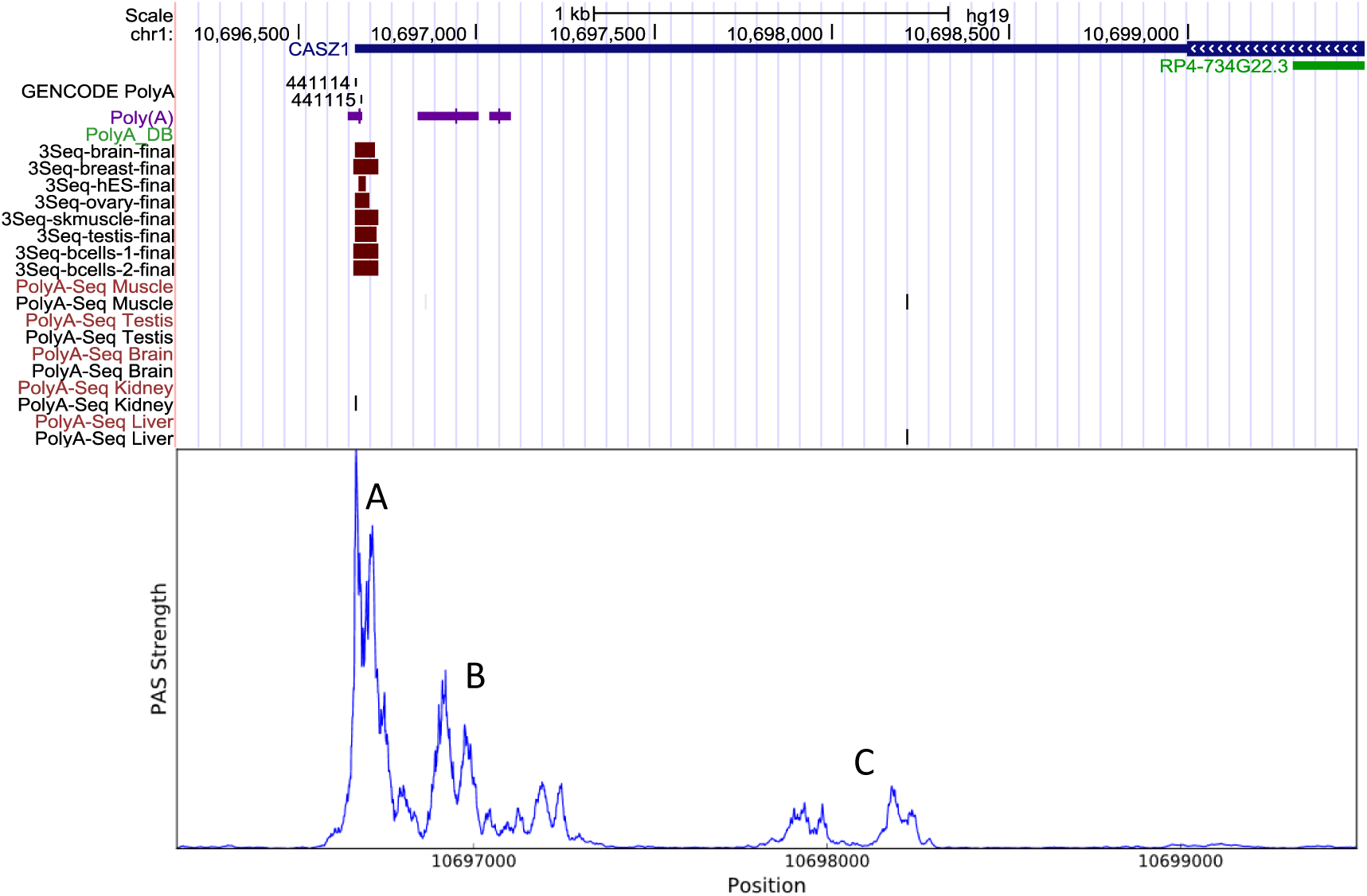
Sample Predicted Polyadenylation Track. Example application of scanning the Conv-Net polyadenylation code across a section of the human genome to identify potential polyadenylation sites. (Top) Snapshot from the UCSC genome browser, showing tracks from top to bottom: GENCODE gene annotations, GENCODE Poly(A) track, predicted and reported PAS from polyA_DB (7, 52), 3’-Seq (22), and PolyA-Seq (fwd. and rev. strands) (40). (Bottom) Predictions from the polyadenylation code.

**S4.**
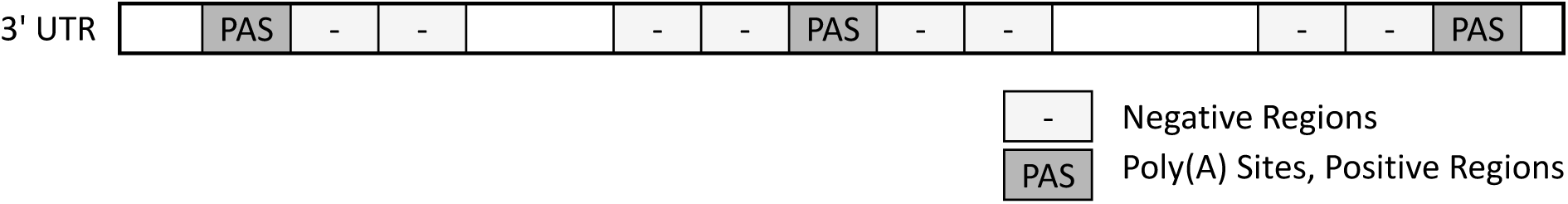
Definition of Positive and Negative Regions for PAS Discovery Evaluation. Two regions immediately adjacent to each PAS are defined as negatives for classification. This ensures that the negatives have similar nucleotide composition compared to the positive sequences. Regions that are not between existing PAS are excluded to avoid including terminal exonic regions. If the spacing between adjacent PAS cannot fit four negative regions, they are also excluded from the negative set.

**S5.**
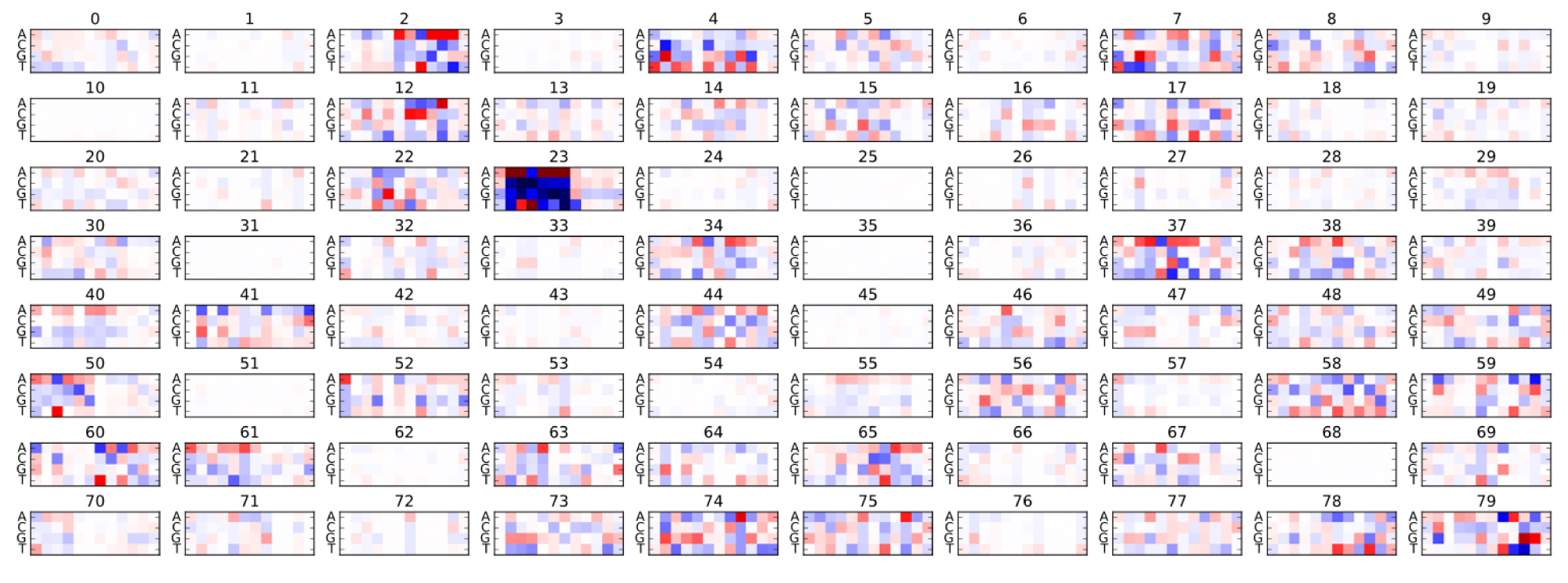
Example Filters Learned by the Convolutional Neural Network. An example set of the 80 filters that are learned by the Conv-Net. All filter has been mean-subtracted and plotted with the same scale (i.e. the max and min for each filter plot is the same). Red and blue denote positive and negative values respectively. Various filters are blank, suggesting the number of filters in the Conv-Net model can be reduced. A filter that detects the two most common polyadenylation signal motifs, ATTAAA and AATAAA can be seen in filter #23. Filters resembling GU-rich elements, such as filter #4 can also be found.

**S6.**
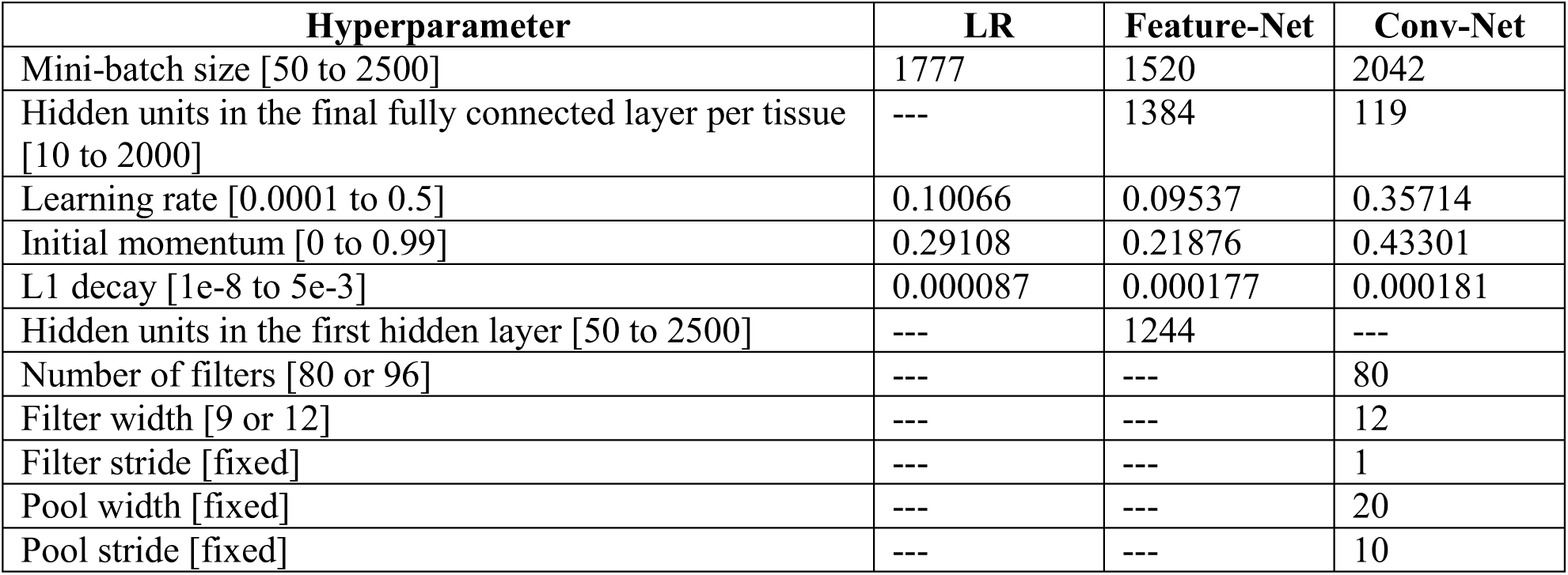
Model Hyperparameters. The following hyperparameters are determined by random sampling and selecting the set that provide the best validation performance. The range each hyperparameter is sampled from is indicated.

